# Integration of multi-omics data shows downregulation of mismatch repair, purin, and tublin pathways in AR-negative triple-negative chemotherapy-resistant breast tumors

**DOI:** 10.1101/2022.05.16.492190

**Authors:** Xiaojia Tang, Kevin J. Thompson, Krishna R. Kalari, Jason P. Sinnwell, Vera J. Suman, Peter T. Vedell, Sarah A. McLaughlin, Donald W. Northfelt, Alvaro Moreno Aspitia, Richard J. Gray, Jodi Carter, Richard Weinshilboum, Liewei Wang, Judy C. Boughey, Matthew P. Goetz

**Author notes:** Contribute equally to the manuscript (Xiaojia Tang, Kevin J. Thompson, Krishna R. Kalari).

## Abstract

Triple negative breast cancer (TNBC) patients who fail to achieve a pathological complete response to neoadjuvant chemotherapy (NAC) will likely experience recurrence of the disease within 3-4 years. Prognostic assessment of early recurrence could facilitate clinical decisions and impact disease survival. This study investigated pre- and post-NAC multi-omics data from both an in-house and a public clinical study to identify biomarkers of NAC response. We observed significant transcriptional differences associated with response in the residual disease (post-NAC biopsies), which did not exist in the pre-NAC biopsies. We further refined the post-NAC transcriptional changes to a 17-gene signature and machine learning models were applied to evaluate the signature’s diagnostic potential (area under the curve ≥ 0.8). Interestingly, the signature was enriched with down-regulated immune genes and several of these genes demonstrated prognostic potential in basal TNBC tumors. Our 17-gene signature provides additional insight into TNBC recurrence, which warrants further investigation.

## INTRODUCTION

Global cancer statistics 2020 (GLOBOCAN 2020) reported breast cancer as the most commonly diagnosed cancer in women, with an estimated 2.3 million new cases per year ^1^. The molecular classification of breast cancer is currently based on the human epidermal growth factor receptor-2 (HER2) gene amplification and expression of sex hormone, estrogen, and progesterone receptors. Triple-negative breast cancer (TNBC) is the most aggressive breast cancer subtype; it lacks estrogen receptor (ER) expression, progesterone receptor (PR) expression, and HER2 gene amplification. The TNBC subtype accounts for approximately 15% of invasive breast cancers. Patients with TNBC have the highest mortality rates compared to the other breast cancer subtypes and are standardly treated with chemotherapy prior to surgery, known as neoadjuvant chemotherapy (NAC). NAC allows a reduction of the size of the tumor burden and decreases the likelihood of nodal involvement before surgery. Most importantly, especially in TNBC, response to NAC is associated with long-term patient outcome^2^. However, nearly 50% of the patients with TNBC have residual disease after NAC^3^. Of those, at least 20-30% of TNBC patients develop an early disease progression within three years and exhibit high metastatic recurrence rates and poor long-term outcome^2,4^.

With the advances in high-throughput technologies, large-scale breast cancer genomics and transcriptomics data have been collected and analyzed to identify biomarkers associated with prognosis as well as treatment resistance ^2,5–12^. The majority of the TNBC studies focused on subtyping or developing signatures associated with prognosis or disease recurrence, mainly utilizing pre-treatment gene expression data from tumor samples. Few studies have evaluated transcriptomics data from TNBC tumors pre-NAC and post-NAC ^2,9–12^. These studies have examined changes in gene expression post-NAC to understand the molecular underpinnings driving treatment resistance in TNBCs. Therefore, identifying genes involved with chemotherapy-resistant disease (residual disease post-NAC) could lead to a better stratification of patients and better therapeutic strategies for TNBC. Balko et al. measured gene expression for 450 transcripts using the NanoString platform from the pre and post-NAC breast tumor tissues and identified the DUSP4 deficiency as an important mechanism of TNBC drug resistance^11^. The Magbanua, *et al*. study (I-SPY1 breast cancer clinical trial) has shown the association between recurrence and response to cell cycle and immune gene sets^8^. Hancock et al. generated pre-and-post transcriptomics data using the Ion torrent platform. They identified increased expression of SMAD2, loss of TP53, and MYC-driven amplification association with chemorefractory TNBC tumors^2^. Interestingly, both I-SPY1 and Hancock et al.’s studies observed the depletion of the immune microenvironment and upregulation of markers related to stemness in the chemoresistant tumors^2,9^.

In this manuscript, we report the results of our interrogation of transcriptome sequencing data from treatment-resistant TNBC tumors. The gene expression data was obtained from a prospective study^3^ of patients with TNBC. Since TNBC represents a heterogeneous group of TNBC subtypes, we focused on androgen receptor (AR) negative TNBC (i.e., ER-, PR-, HER2- and AR-), given that a paucity of targeted therapies exists for patients with AR-negative TNBC disease. We first developed a signature associated with early recurrence using the post-NAC gene expression data from chemoresistant TNBC tumors. That signature was then compared with the patient’s pre-NAC transcriptomics, whole-exome sequencing (WES), and Reverse Phase Protein Array (RPPA) data. Moreover, we then validated the signature using I-SPY1 pre-and-post-NAC TNBC data. We then verified and strengthened our results with publically available TNBC molecular datasets. We finally identified a 17-gene signature that can predict post-NAC patients with a high risk of recurrence.

## RESULTS

From the prospective BEAUTY neoadjuvant study, we analyzed 18 pairs of triple-negative breast tumors with RNA-Seq data from pre-NAC and post-NAC after surgery (patients who failed to respond to paclitaxel and anthracycline/cyclophosphamide combination therapy). Based on our investigation and existing studies, it is known that the luminal androgen receptor (LAR) TNBCs are biologically and clinically distinct from non-LAR tumors. Hence, we removed the three TNBC LAR tumor pairs identified by LAR-Sig from further analysis. In addition, tumor bed analysis was performed using an *in silica* deconvolution method, indicating that three post-NAC biopsies contained low epithelial composition, both luminal and basal. Hence, we removed those three also from the analysis. We finally investigated 12 patients with non-LAR TNBC tumors with matched pre and post-NAC gene expression data. Of these 12 patients, four had an early recurrence (less than two years after completion of NAC), and eight remained recurrence-free for more than four years. Furthermore, for these 12 non-LAR TNBC patients, all patients had pre-NAC WES data, and 9/12 had paired protein and phoshoprotein (RPPA) data.

### Post-NAC transcriptomics analysis showed clusters of genes differentially expressed on chr1q and chr17q

We investigated post-NAC RNA-Seq data for transcriptional differences associated with the response after standard chemotherapy treatment. We evaluated 20,543 genes assayed from both pre- and post-NAC biopsies. A co-inertia plot of paired samples is presented in **Figure 1A**, using the first two principal components. In the plot, we noted a slight trend of separation between patients with early-recurrence of disease (ERC) and non-recurrence (NRC) for the first principal component. This component explains 20.3% of the variance observed in the pooled dataset.

**Figure 1:**
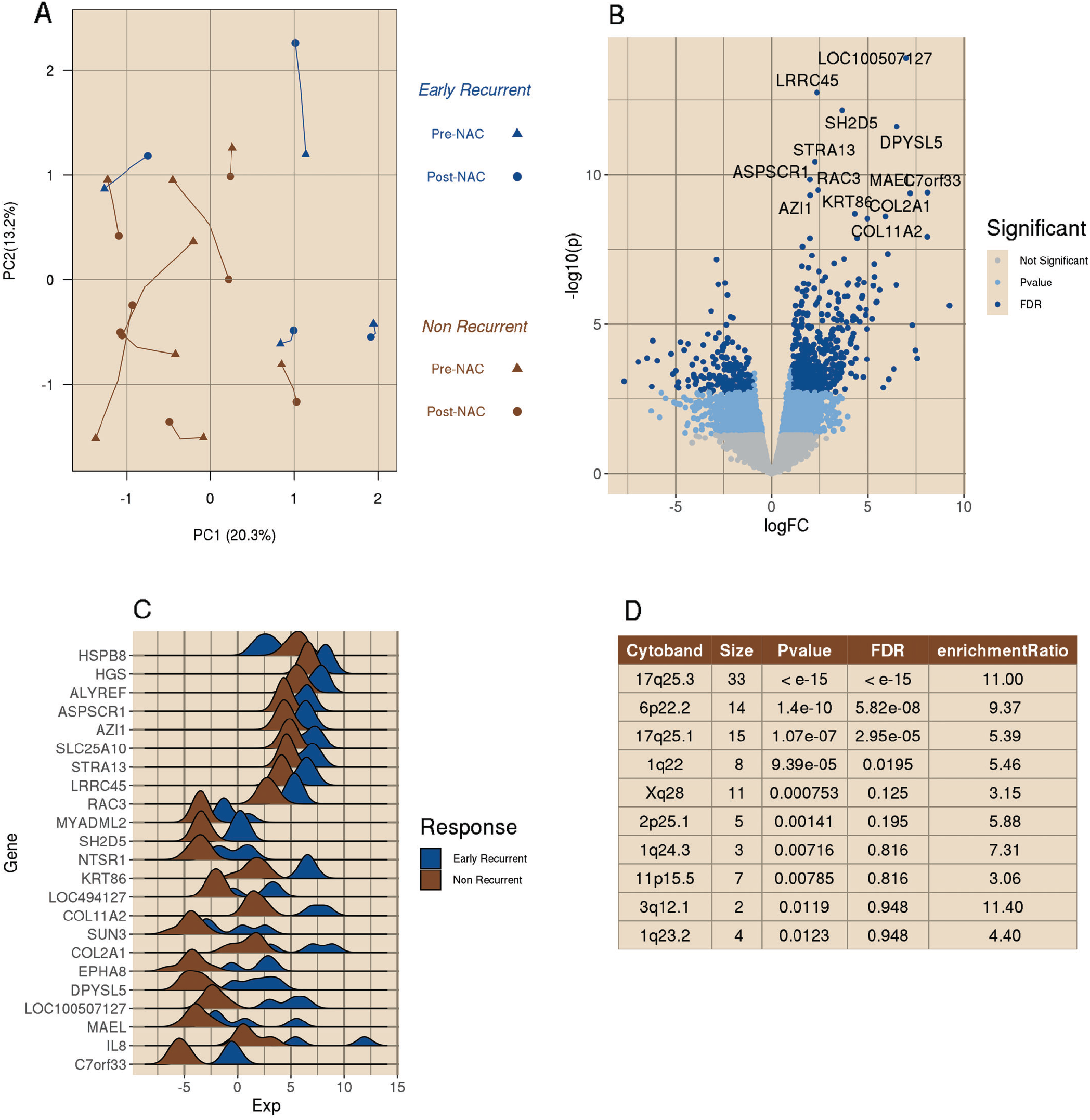
Post-NAC upregulation in early recurrence. A) A co-inertia plot of the top two principal components from the paired bulk RNA-Seq dataset for 12 BEAUTY TNBC non-responders. Lines between points connect the paired samples, with pre-NAC visit indicated with the triangular point and post-NAC resection as points. The four Early Recurrent are presented in blue, and the eight Non-Recurrent patients are presented in brown. We observe a slight trend for non-recurrent to shift left on PC1 and early recurrent shifting away from the center on PC2. B) (top right) A volcano plot of the bulk RNA-Seq DE compares post-NAC surgical samples of recurrent versus non-recurrent. A total of 20,543 genes were tested, with 2,944 (14.3%,) having a p-value <.05 and an absolute logFC greater than 1 (indicated with in the light blue), with the majority (1,859, 63.1%) of those being upregulated in the early recurrent samples. There were 660 genes that were significant after correcting for the false discovery rate, indicated with the darker blue. Gene symbol labels indicate the top 13 genes meeting the significance threshold of p-value < 1e8 and FDR < 1e-5, all of which are upregulated in the early recurrent samples. C) Ridges plot of the twenty-three genes with FDR < 1e-4, distributions for the early recurrent tumors are indicated in the dark blue, and the distribution of the non-recurrent samples presented in brown. D) Presents the results table for the top 10 cytobands enriched in DE genes between early recurrent and non-recurrent TNBC samples.

We first compared the pre-NAC transcriptome gene expression between ERC and NRC groups. We identified only 11 genes to be differentially expressed after correcting for multiple testing (FDR<0.05). Conversely, we evaluated ERC and NRC tumors in the post-NAC gene expression data. We identified 660 genes differentially expressed with at least 2-fold change and FDR < 0.05 (**Figure 1B, Supplementary Table 1**). Among these genes, 478 (72.4%) were upregulated in ERC tumors (**Figure 1B**). The list of the top 23 significant genes based upon an FDR < 1e-4 is shown in **Figure 1C.**

Further over representation analysis (ORA) identified 10 cytobands that were significantly enriched in the 660 DE genes (**Figure 1D**), including 17q (17q25.3, 33 genes, p-value less than 10^-15^ and 17q25.1, 15 genes, p-value 1.01×10^-1^), 6p (6p22.2, 14 genes, p-value 1.40×10^-10^), 10q (10q22.3, 9 genes, p-value 6.51×10^-5^), and 1q (1q, 8 genes, p-value 9.39×10^-5^; 1q24.3, 3 genes, p-value 0.007; and 1q23.2, 4 genes, p-value 0.01).

### Genomic analysis of pre-NAC data confirms amplifications in chromosomes 1q and 17q in the early-recurrence group (ERC) compared to the non-recurrent (NRC) group

Since WES data of post-NAC tumors were unavailable, we examined copy number alterations in the pre-NAC patient tumors between ERC and NRC groups. We identified a common gain/amplification in a large region of the chr1q (q21.3-q24.2,395 altered genes) and chr17q (q25.3, 97 altered genes) in at least 3 out of 4 recurrent patients; however, they were not detected in any of the eight non-recurrent patients (p-value <=0.012 by Fisher’s exact test). The 1q and 17q gain events involved 492 genes (**Supplementary Table 2**). Similarly, we also identified a deletion of 4 genes (in chr4p) among the ERC group. A circos plot depicting the altered copy number regions is presented in **Figure 2A**. We also present histogram views of the copy gains in the early-recurrent samples for the 17q and 1q regions along with the elevated expression levels observed with these copy gains using pre-NAC and post-NAC RNA-Seq data (**Figure 2B and 2C,** respectively). Below the circos plot, we present the chromosome 4p15.2-15.32 region, where we detected an increased copy loss in the early recurrent patients. Again, a decrease in gene expression was observed across the region (**Figure 2D)**.

**Figure 2:**
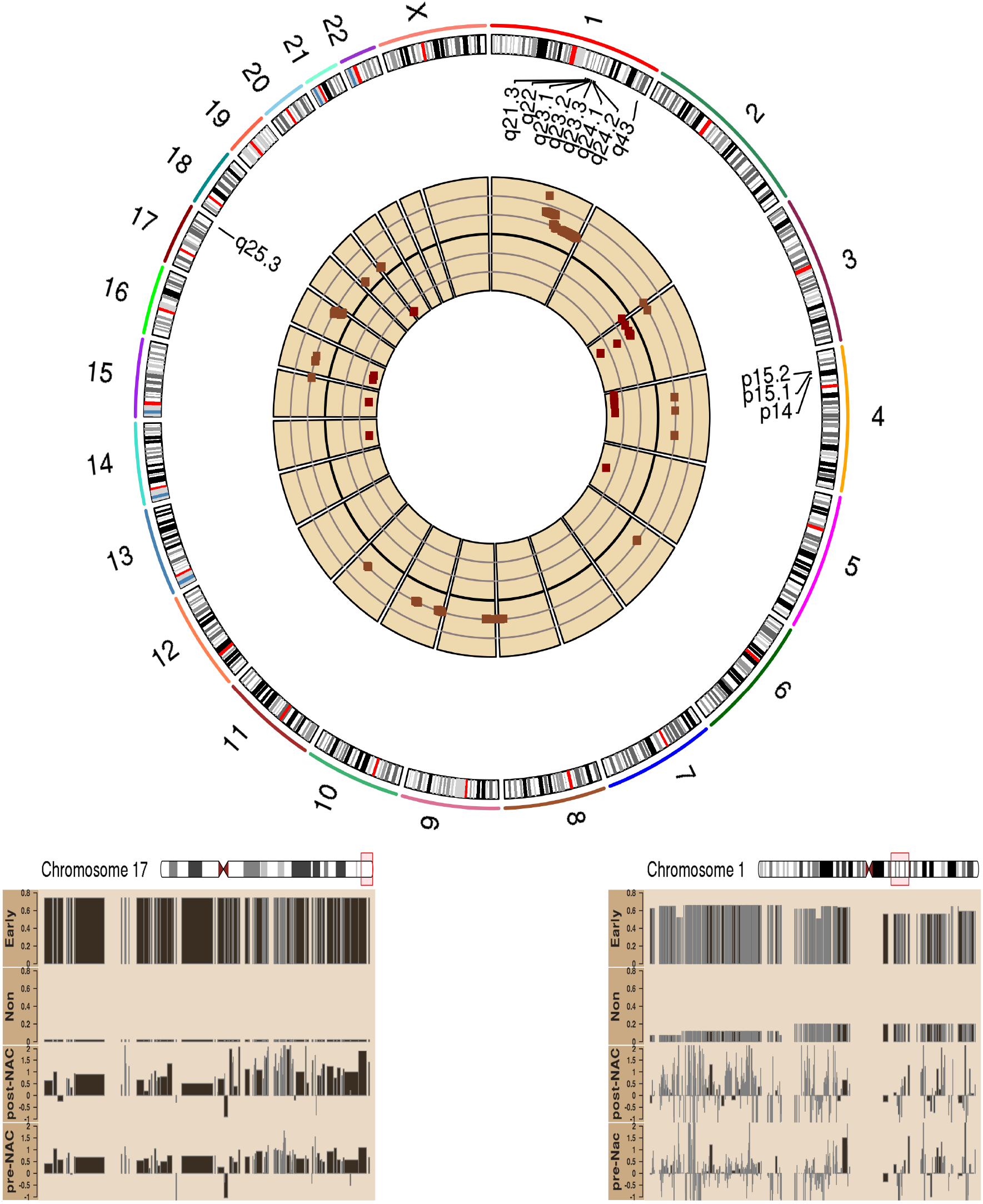
Circos plot of the gene-level copy number alteration. Gene-level CNA events with a p-value of less than 0.05. Cytoband regions that were observed to be over-represented with genomic alterations are labeled, including 17q25, 1q24-24, and 4p15.2-.32. The hazard ratio of the copy gains is presented in the outer track (lighter brown), and the hazard ratio of copy loss is shown in the inner circle (darker brown). Gain on chr1q and chr17q were observed in 3 out of 4 recurrent patient tumors and the regions are expanded, respectively top to bottom, on the right. Elevated expression levels are observed, which coincide with these copy gains. Similarly, decreased expression is observed in the 4p15.2-4p15.32 region, coinciding with the observed copy loss.

### Somatic Mutation showed frequent mutations in TP53 and MUC4

We investigated the somatic mutations in the pre-NAC sample biopsies using WES tumor and blood DNA data. We identified 392 high/moderate impact somatic mutations corresponding to 366 genes in the four early-recurrent patient tumors and 545 high/moderate impact mutations in 519 genes in the eight non-recurrent patient tumors, respectively (**Supplementary Table 3**). Due to the small sample size, low mutation frequency across genes and low overlap among the genes were observed between the ERC and NRC groups. It was noted that the most frequently mutated genes across our cohort were TP53 (all four ERC samples and five of eight NRC samples) and MUC4 (two of the four ERC samples and three of eight NRC samples), consistent with the previous publication^13^.

### RPPA analysis indicates pre-NAC proteomic differences

We evaluated protein expression (232 proteins) and phosphorylation (63 phosphorylation sites) data obtained for the pre-NAC and post-NAC tumors using RPPA. The pre-NAC RPPA data were compared between the ERC and NRC groups. We observed that 30 proteins were significantly different (p-value < 0.05), including ten phosphorylation sites (10/63, 15.9%) and 20 total proteins (20/232,8.6%) among the pre-NAC samples. In contrast, there were only four protein level variations and no phosphorylation site differences in the post-NAC RPPA data. See **Supplementary Tables 4-5** for full results.

### Down-regulation of Tubulin and upregulation of growth receptors in the non-recurrent post-NAC tumors

A parallel single-sample gene set analysis was performed in an attempt to gain more insight from the gene expression data and resolve the challenges of a small sample size study. We performed Gene Set Variation Analysis (GSVA) and obtained individual gene set scores for all non-LAR TNBC tumors with the pre-NAC and post-NAC RNA-Seq data. We evaluated three comparisons (post-NAC difference between ERC vs. NRC, pre and post-NAC (paired) ERC, and paired NRC). Among the total 2282 gene sets, we identified 251 gene sets (**Supplementary Table 6**) significantly altered comparing the ERC and NRC group in the post-NAC tumors (p-value < 0.05).

In addition, we performed a hierarchical clustering analysis of log fold changes for 251 gene sets, **Figure 3**. The log fold changes represented the change in post-NAC data and the difference between the tumors for both the ERC and NRC groups. The average silhouette width, a metric of within-class similarity and between-class dissimilarity, was 0.90 for two gene set clusters and was markedly reduced for three gene set clusters (0.61). We observed that the first cluster of 61 gene sets were downregulated in the post-NAC non-recurrent patients, compared to the post-NAC early recurrent group as well as the own pre-NAC biopsies. Tubulin (TUBB4A) was the most frequent gene in these gene sets, present in at least eleven (18.0%) gene sets, whereas the pre-NAC expression levels for tubulin were not significantly different. Conversely, the second cluster contains 190 gene sets that were upregulated in the post-NAC non-recurrent tumors. Furthermore, this cluster was down-regulated in the post-NAC early-recurrence cohort compared to their pre-NAC ERC sample biopsies. The heatmap image is presented with the gene clusters labeled, and the most commonly represented genes in the gene sets constitute the word clouds in **Figure 3**. In summary, cluster 1 gene sets that were downregulated in the post-NAC NRC patients were predominately associated with tubulin, and in cluster2, the gene sets that were upregulated include growth receptors such as EGFR, TGFBR2, and GHR, as well as tumor suppressor genes found in apoptotic gene sets.

**Figure 3:**
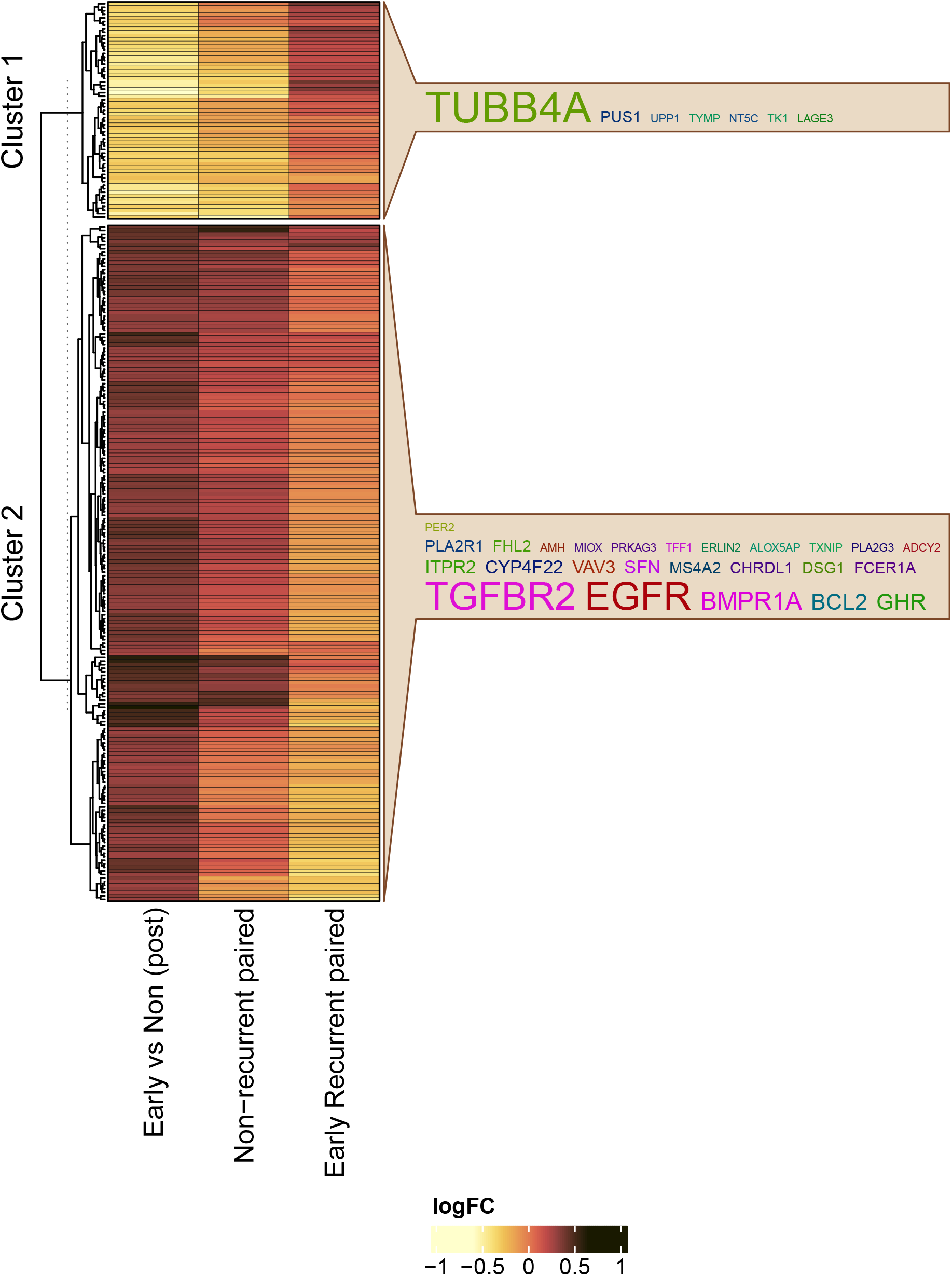
GSVA analysis showed pathway-level/gene set activity differences between the early recurrent and non-recurrent groups. In the figure, column 1 - Non-recurrent paired pre-NAC vs. post-NAC), column 2 – Early recurrent paired pre-NAC vs. post-NAC, and column 3-post-NAC (early recurrence vs. non-recurrence). Unsupervised clustering showed two clusters with a distinctive pattern between the two groups (heatmap panel). Bottom) Cluster 1 gene sets that were upregulated in the early recurrent group included metastasis-promoting gene sets, DNA mismatch repair, and tp53 gene sets, and the most commonly observed gene was TUBB4A. Cluster 2 included gene sets like FOXO, TGF-beta, and apoptotic gene sets known to be tumor suppressors.

### I-SPY1 confirmation of the core gene set signature from the BEAUTY TNBC cohort

We investigated publically available data from the I-SPY1 study to validate our pathway analysis findings. Due to the platform differences between the I-SPY1 (microarray) and BEAUTY (RNA-Seq) datasets, we calculated the gene set scores for 188/251 (74.9%) gene sets using the GSVA method. Of the 188 gene sets, 56 (29.8%) gene sets demonstrated high similarity between the I-SPY1 and BEAUTY studies (**Supplementary Table 7** of 56 pathway). Among these 56 gene sets, 17 gene sets (cluster 1, **Figure 3**) were down-regulated, and 39 (cluster 2, **Figure 3**) were upregulated in the post-NAC I-SPY1 early-recurrence tumors. Cluster 1 could not recapitulate the same set of genes as TUBB4A was not assayed on the microarray platform used for the ISPY trial. We observed five consistent genes in cluster 2, including ERLIN2, FCER1A, TGFBR2, PER2, and EGFR. These genes are common to gene sets, including MYC targets, TGFβ, FOXO, and PI3K gene sets.

### Identification of 17-gene signature to predict recurrence of cancer in chemoresistant tumors

We identified 113 differentially expressed genes either up/down-regulated in the same direction when comparing the ERC and NRC patients in the I-SPY1 and BEAUTY studies related to 56 gene sets (listed in **Supplementary Table 7**). We assessed the prognostic value of these 113 genes in a publically available TNBC cohort (*n*=392 independent dataset pooled from several TNBC studies) using the Kaplan-Meier (KM) plotter online breast cancer database. Our analysis shows that 17 genes were significantly associated (log-rank test p-value < 0.05) with relapse-free survival (RFS) in the TNBC cohort (**Figure 4, Supplementary Figure 1**). Further investigation into this same subcohort with lymph nodes positive disease (N=189) showed that 10 out of the 17 genes still remained significant (Table 1).

**Figure 4.**
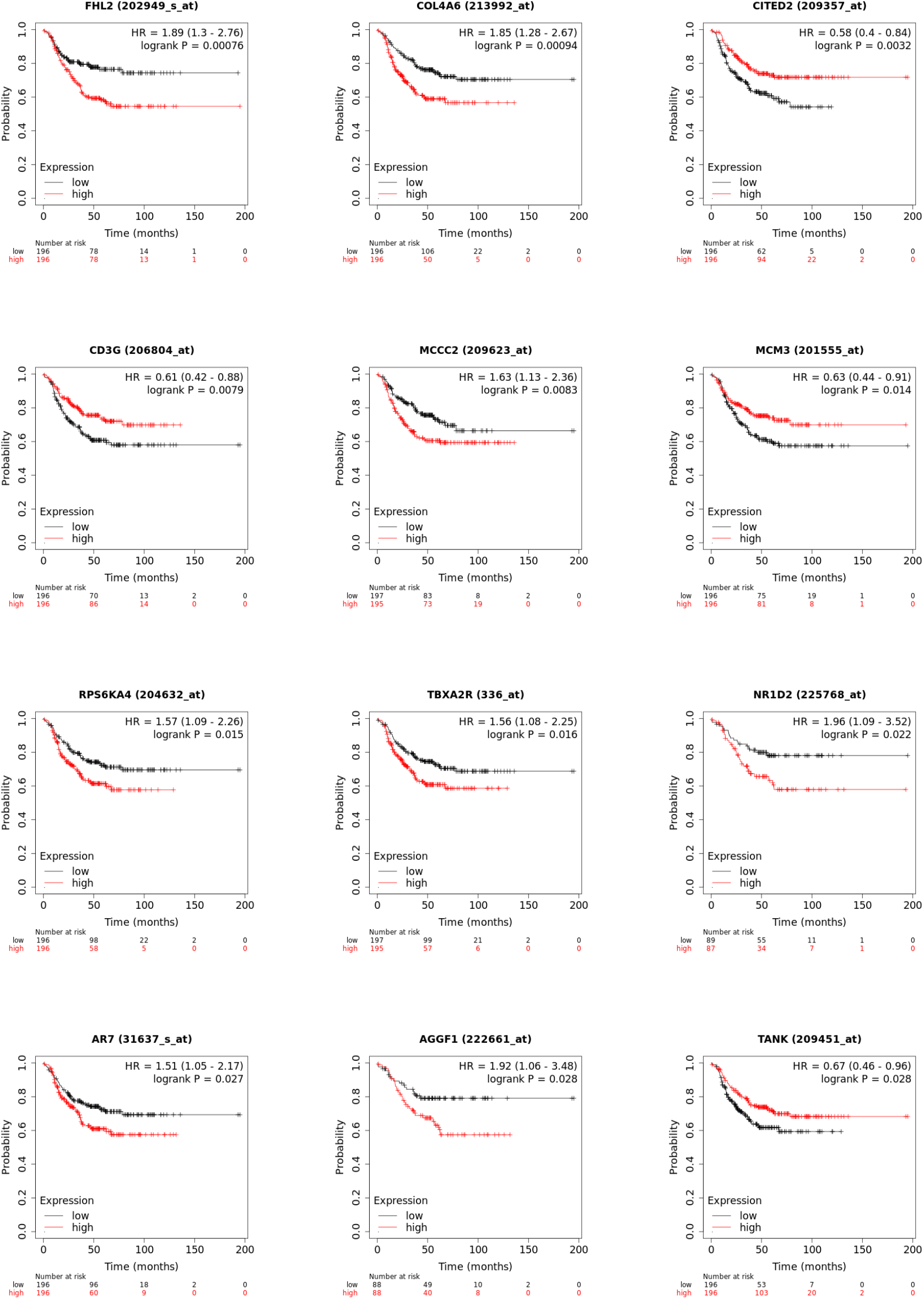
Seventeen genes were associated with survival analysis in an independent TNBC cohort (N=392) from the KM plotter database. Sixteen of them are presented in this plot.

**Table 1.**
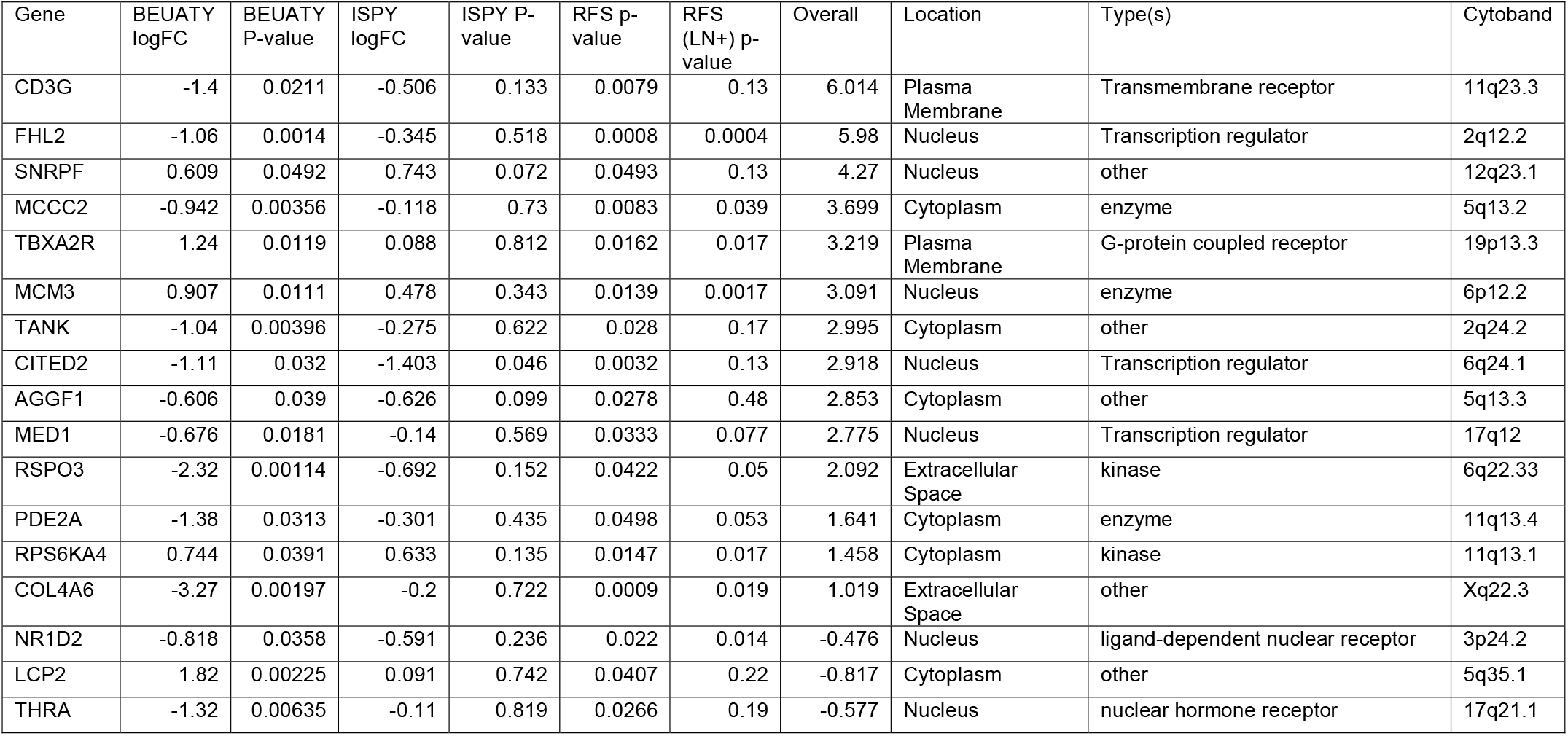
List of the 17 gene signature.

### Prediction of recurrence using chemoresistant gene expression data, machine learning models, and cross-validation methods with the 17 gene signature

Since there are no independent neoadjuvant breast cancer studies with publically available data to validate our signature, transcriptomic data from the BEAUTY and I-SPY1 studies were pooled together to evaluate the 17-gene signature using machine learning models. We examined six classification algorithms, including a generalized linear model (glmnet), k nearest neighbors (k-NN), neural networks, random forests, support vector machine (SVM), and kernel SVM. The models were implemented as wrappers in the R package superlearner using gene down selection methods (Pearson’s correlation, Spearman’s rank correlation, and a random forest). Given the lack of sampling power, we implemented a three-fold cross-validation strategy, which we believe provided the most reasonable cross-validation strategy without subsampling the data into too trivial of a sample representation. We monitored the area under the receiver operating curve as indicative of the model’s classification performance, provided in **Figure 5**. The rank-based correlation (dark brown diamond) consistently underperformed other feature selection approaches (**Supplementary Figure 2**). Most importantly, the ensemble of seventeen (All in **Figure 5**, tan diamond) genes was consistently the best performing approach. We present the distribution of AUCs observed for each classification wrapper, **Figure 5**. The models are ordered by increasing median AUC values, and kernel support vector machines, k nearest-neighbor (kNN), and random forest were the best performing modeling approaches, with the ensemble of all 17 genes achieving AUCs of 0.84, 0.88, and 0.88, respectively.

**Figure 5:**
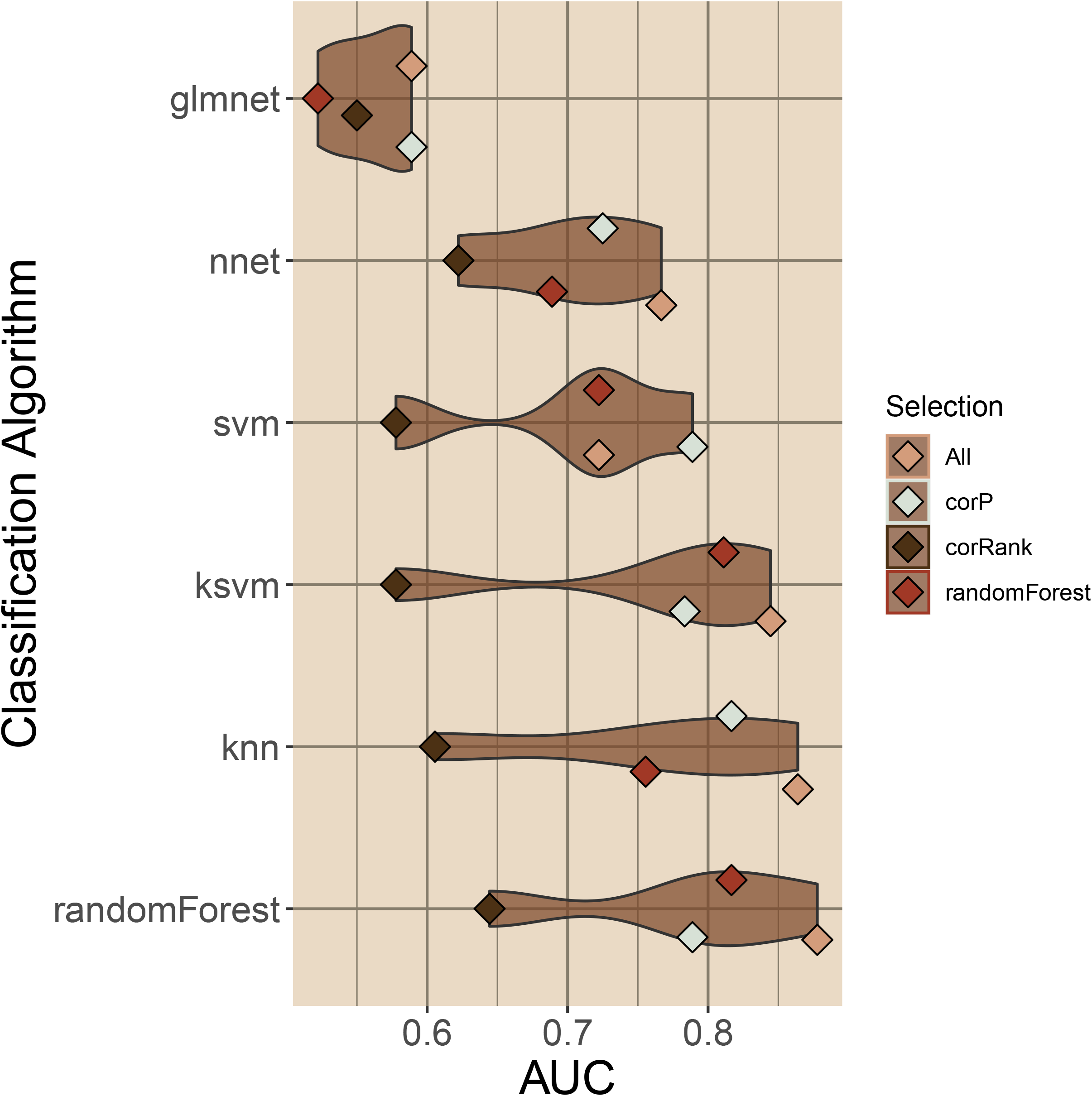
Classification assessment of seventeen genes associated with triple-negative breast cancer recurrence in chemoresistant tumors. Six classifiers were evaluated with the SuperLearner package employing seventeen and three down selection methods. The mean cross-validation AUC is plotted as a point for each classification method, and the four feature selection methods used are shown. We observed that three models, kernel support vector machines, kNN, and random forest achieved the best overall median classification AUC.

### Differential expression of 17 genes in post-NAC TNBC ERC and NRC cohorts compared to paired pre-NAC tumors, TCGA tumors, and TCGA normal-adjacent tumors and association with pCR

The expression profiles of the 17 genes were evaluated in 1) post-NAC and pre-NAC ERC vs. NRC cohorts, 2) TCGA paired tumor and normal-adjacent TNBC samples^14^, 3) paired non-LAR TNBC tumor vs. normal adjacent TCGA samples. As shown in **Figure 6**, we verified that the differential expression of these 17 genes was specific to post-NAC biopsies; none of the genes were observed to be differentially expressed among the pre-NAC biopsies. Further, we evaluated the expression among eleven paired TNBC tumors and normal adjacent samples from TCGA and observed that the expression profile was consistent among the tumor samples, with 52.9% (nine of the seventeen) surveyed being differentially expressed. Additionally, this expression pattern was confirmed among the nine paired non-LAR TCGA TNBC tumors and normal-adjacent samples (**Supplementary Figures 3 & 4, Figure 6**). Moreover, we evaluated whether the expression of these 17 genes was specific to non-LAR compared to LAR TNBC samples using an independent cohort (N=1209). We observed 13 of the 17 (76.5%) genes differentially expressed (**Supplementary Figures 5**) between the LAR and non-LAR TNBCs. The cohort of 1,209 TNBC (pre-NAC) samples included Affymetrix microarray data along with RNA-Seq data, and the gene RSPO3 was not assayed in the microarray platform. This combined cohort was scaled together, and while the fold change of the genes appeared relatively small, the observed p-value represented the most significant differences. Finally, among the combined cohort, there were 878 non-LAR TNBCs (432 with pCR and 446 with residual disease) who had pathological complete response data. We observed that only 5 (31.3%) of the 16 genes (RSPO3 was not assayed) were differentially expressed (**Supplementary Figures 6**). In summary, the 17 genes identified in this study are associated explicitly with the prediction of post-NAC recurrence in non-LAR TNBCs with residual disease after NAC (Supplementary Figures 7).

**Figure 6:**
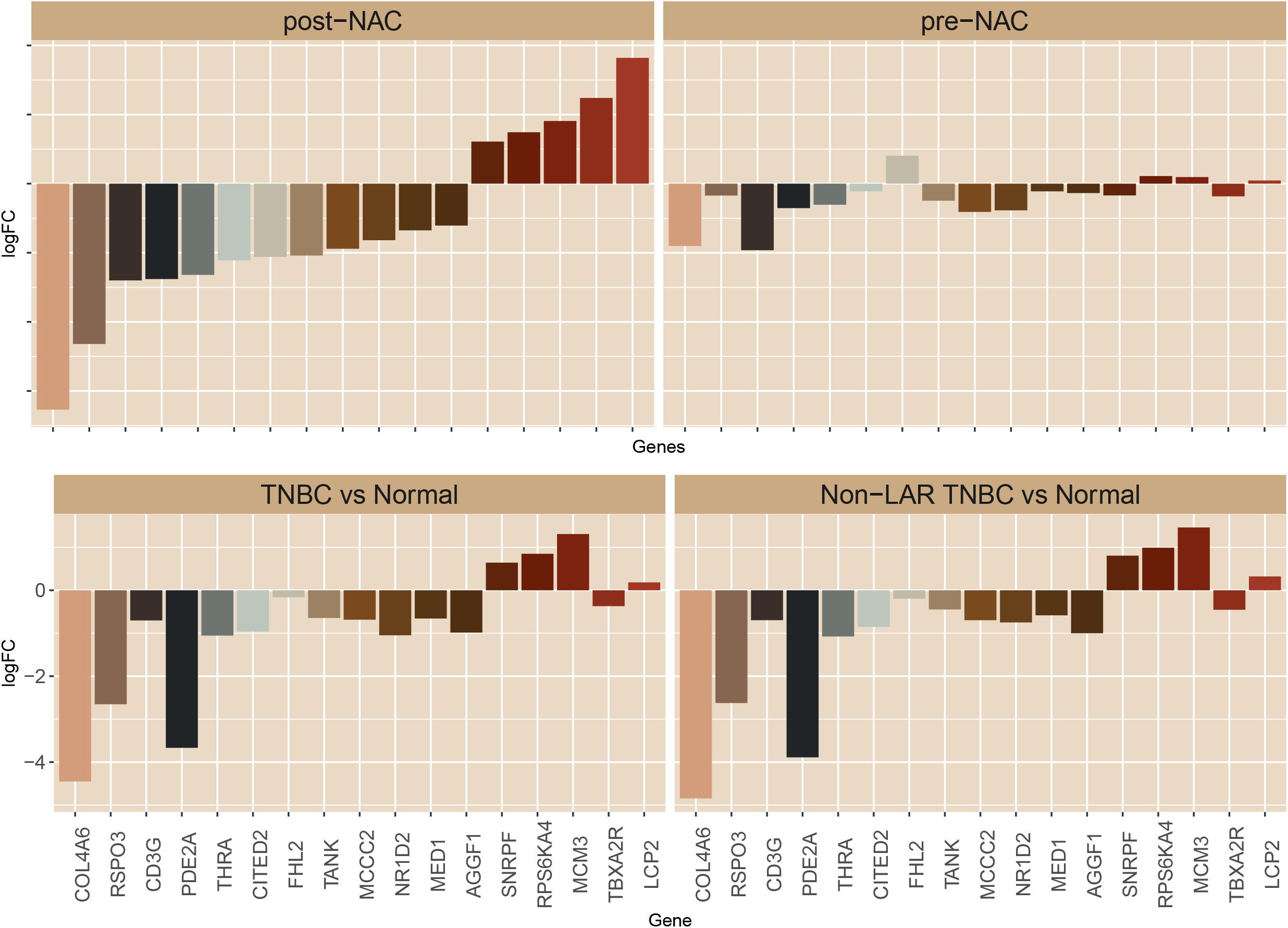
Expression profile of seventeen, which distinguishes early recurrence in post-NAC biopsies. The seventeen were observed to be differentially expressed in TNBC post-NAC biopsies (top row left). However, none of these genes were differentially expressed in the pre-NAc biopsies (top row right). Four of the genes (FHL2, TANK, PDE2A, and RSPO3) are related to immune response signaling, and they were all observed to be down-regulated in the early recurrent samples. The expression profile was confirmed to arise from tumor tissues (bottom row) compared to paired normal-adjacent tissues. Moreover, the expression profile was specific to non-LAR tumors (Supplementary Figures 3-6).

## Discussion

Triple-negative breast cancers are a phenotypically heterogeneous disease that presents in a younger population of women. The TNBCs currently lack therapeutic targets, and neoadjuvant chemotherapy is the standard of care from stage I to stage III diagnosis. The 3-year survival rate of TNBC patients who achieve pathological complete response is 94%. However, the 3-year survival rate plunges to 68% for those who fail to achieve pCR, and approximately 50% will develop recurrent disease within the first 3-4 years^2^. Hence, we investigated the chemoresistant BEAUTY TNBC tumors for potential biomarkers delineating early recurrence patient’s landscape from non-recurrence profiles.

First, we investigated the differential expression at pre-NAC and observed a reduced set of differentially expressed gene features compared to that of the post-NAC. This suggests that the pre-NAC tumors represent multiple steady-states rather than a simple two-state model. The implication would be that post-NAC biopsies demonstrate a two-state model convergence based upon selective therapeutic pressure. Among the 660 differentially expressed genes (FDR < 0.05 and |logFC|>1), we observed enrichment of genes in post-NAC data in a few cytoband regions (**Figure 1**). Two of these regions (17q25 and 1q23-24) were confirmed to demonstrate copy gains in the pre-NAC biopsy, which coincided with the elevated expression observed in the post-NAC early recurrent samples. Three genes found in the 17q25 region are involved in sumoylation: NUP85, CBX2, and CBX4. Similarly, the cytoband 4p15 presented with a copy loss in the early recurrent tumors (Pre-NAC), and the post-NAC also demonstrated decreased expression (**Figure 2**).

A parallel track of analysis using topological gene set measurements was adapted to navigate the challenges we faced, given the relatively low sampling power. We conducted a GSVA analysis on the post-NAC BEAUTY gene expression data and leveraged our observations with the independent I-SPY1 studies post-NAC microarray data. We observed that 251 gene sets were altered between the early recurrent and non-recurrent BEAUTY TNBC tumors(**Figure 3)**. Cluster analysis of the 251 gene sets identified two clusters, and the first cluster includes TUBB4A, the therapeutic target of paclitaxel. The gene sets were upregulated in the early recurrent group and included metastasis-promoting gene sets, DNA mismatch repair, and TP53 gene sets. In contrast, the second cluster of gene sets were down-regulated in the early recurrent samples involving tumor suppressor gene sets, including FOXO signaling, TGF-beta signaling, and apoptosis (**Figure 3**).

Due to the technology differences between the microarray (I-SPY1) and RNA-Seq (BEAUTY) data, only 188/251 gene sets (gene sets) from the BEAUTY study were also investigated in the I-SPY1 study. We confirmed 56 gene sets in the I-SPY1 data to be significant and concordantly altered. Further, the 56 co-observed significant gene sets were represented by 113 genes that were concordant in direction and differentially expressed in the two datasets. We investigated 113 genes further in an independent cohort of TNBC patients (N=392) using the gene expression data and the online tool KM-plotter. We identified seventeen genes associated with recurrence-free survival in 392 TNBC samples (**Figure 4**). Later, we conducted a machine-learning analysis of these genes and demonstrated a robust ability to predict recurrence using chemoresistant gene expression data. Six separate classification algorithms were explored, with random forest, kNN, and kernel SVM being the best performing algorithms, using 17 genes indicating that no individual gene was a dominant classifier (**Figure 5**). Differential expression analysis of 17 genes in post-NAC TNBC early recurrent and non-recurrent cohorts was significantly different from pre-NAC tumors (**Figure 6**). The 17 genes were also differentially expressed in the paired TCGA and normal-adjacent tumors. In an independent cohort of 1209 TNBCs, we found that the genes are differentially expressed in non-LAR TNBCs compared to LAR-TNBCs, and only five genes in non-LAR TNBCs were associated with pCR (**Supplementary Figure 4**).

In summary, we confirmed that the expression of these 17 signature genes is derived from non-LAR TNBC tumors and that expression differences were specific to post-NAC treatment. Four genes (FHL2, TANK, PDE2A, and RSPO3) were well documented to be associated with breast cancer and were observed to be down-regulated in the post-NAC early recurrent samples. PDE2A is a phosphodiesterase that regulates mitochondrial respiration and mitogenic clearance^15,16^. FHL2 is a zinc finger transcription factor that has been associated with several cancers, including ovarian and cervical cancers^17–19^. We note that the I-SPY1 trial observed a contradictory increase in interferon signaling associated with shorter recurrence-free survival among non-responding patients^9^. We observed that two molecules, RSPO3, and TANK, are related to NF-κB signaling and survival response through the induction of inflammatory molecules. RSPO3 is a matricellular protein that activates WNT signaling via the canonical and non-canonical gene sets. Its involvement in cell development and growth has implicated its association with colorectal and lung cancers. RSPO3 downregulation has been involved in prostate cancer invasiveness, and it interacts with the inflammatory mediator IL-1β^20,21^. While two SNPs (*rs17705608* and *rs7309*) in the TANK gene have been associated with breast cancer risk^22^, it is involved in TNF-mediated signal transduction. Most importantly, the TANK, the TRAF family member, binds the NEMO (IKKγ) to induce inflammation through the IKK complex and NF-κB signal transduction^23–25^. These findings suggest that initiating inflammation signaling via NF-κB signal transduction is integral to recurrent free disease. Moreover, we observed significant downregulation (−2.91, *p*-value 0.044) of the key apoptotic protein, BCL2, suggesting that the inflammation must be accompanied by immune infiltration and subsequent cell death.

We systematically analyzed the in-house and publicly available TNBC gene expression NAC datasets using computational biology and machine learning methods. We concluded that the post-NAC non-LAR TNBCs change significantly during the treatment compared to the baseline pre-NAC tumors. We have shown that the 17 genes identified in this study are novel biomarkers that predict recurrence in post-NAC residual tumors. Our analysis has also demonstrated that initiating inflammation signaling via NF-κB signal transduction is integral to recurrence-free disease.

## METHODS

### Materials or datasets

#### A chemorefractory cohort from the Mayo Clinic breast cancer study

The Breast Cancer Genome Guided Therapy Study (BEAUTY) is a prospective institutional review board–approved NAC clinical trial (NCT02022202). As previously described^3^, The BEAUTY study enrolled 140 patients with invasive breast cancer, including 42 patients with TNBC. Eighteen of the patients with TNBC had residual disease at surgery. We obtained pre-NAC and post-NAC transcriptomics data for these 18 cases and identified luminal androgen positive (LAR) cases using our in-house algorithm (LAR-Sig). Due to significant clinical, molecular, and pathological differences we previously observed between the LAR and non-LAR tumors, we focused our analysis on the non-LAR (AR negative) TNBC patients^26–30^. We also removed three samples as the post-NAC biopsies contained minimal epithelial tissue (see below). The final cohort for paired visit analysis included four early recurrent and eight nonrecurrent samples. In addition to gene expression from transcriptome RNA-Sequencing, we also investigated copy number alteration and somatic mutation data from whole-exome sequencing (WES, pre-NAC due to availability) from the BEAUTY study. Here we summarize our findings of interrogating paired gene expression data, paired protein expression data (described below), and pre-NAC WES data from the BEAUTY cohort.

#### Reverse Phase Protein Array data from the BEAUTY breast cancer study

Lysates from paired tissue preparations of the paired breast tumor tissue samples at pre-NAC and post-NAC time points from the BEAUTY study were provided for evaluation by Reverse Phase Protein Array (RPPA). Relative protein levels were determined by interpolation using a super curve provided by MD-Anderson Cancer Center for functional proteomics^31^. Of the twelve patients in the early recurrent and non-recurrent groups, eight had RPPA data at pre-NAC, and six had RPPA data at post-NAC visits. We performed two-sample t-tests with pooled variances on the log2 transformed protein levels for 295 antibodies between the pre-NAC and post-NAC data.

#### Tumor bed assessment using digital deconvolution methods

A digital deconvolution was performed on the post-NAC samples using gene expression data to ensure sufficient epithelial cells were obtained rather than tumor beds. Twenty cell types were previously observed in a single-cell RNA sequencing experiment, originating from five patients with primary TNBC^32^. We obtained the single-cell gene expression counts and t-SNE clustering scheme and constructed a balanced dataset with prevalent and low abundance cell types. Our balanced dataset consisted of eighty nearest neighbor cells to each of the twenty centroids to adequately represent the least prevalent cell type (i.e., 106 immature perivascular-like fibroblasts, imPVL). The mean-dropout relationship (high zero counts while maintaining a high mean expression level) was evaluated, and the features were reduced to 3,205 genes (curve fitting parameters of a=1.5 and b=1.1)^33^. A deconvolution model was then constructed using the balanced TNBC single-cell data and CIBERSORTx ^34,35^ method. The deconvolution was implemented with the twenty cell types merged into nine lineages. Each BEAUTY study sample with bulk RNA-Seq and phenotype was individually evaluated with pseudo replication to achieve a sampling set larger than the nine cell types. Three paired RNA-Seq samples (two nonrecurrent and one early recurrent) were removed as tumor bed biopsies.

#### I-SPY1 trial gene expression data

The clinical and molecular data from the I-SPY1 breast cancer trial have been described in detail previously ^36,37^. The normalized expression data were downloaded from the gene expression omnibus (GEO) database with a GEO ID: GSE32603^9^. From the I-SPY1 GEO data set (N=141, with baseline sequencing data), we selected patients with TNBC (N=39, 27.7%) that were resistant to NAC (N=23, 59.0%). We also considered only paired TNBC samples (N=10) having gene expression data at both pre-NAC treatment (T1) and post-NAC or surgery (TS) time points. Further, like the BEAUTY breast cancer study, we also considered the paired TNBC samples (N=5) from patients who had an early recurrence (in less than two years after NAC) and patients who remained non-recurrent for more than four years for gene expression data analysis(N=4).

#### Gene set analysis

Sample-wise pathway analysis of the RNA-Seq data was carried out using the gene set variation analysis (GSVA) method ^38^ for the curated (C2, *n*=2,232) and hallmark (H, *n*=50)gene sets from MSigDB version 7.1M^39^. GSVA scores were calculated for the pre-NAC and post-NAC gene expression datasets. The scores were then evaluated using linear regression with empirical Bayes statistics with limma package^40^. Gene sets were determined to be significant with a p-value < 0.05. Gene sets and individual genes were filtered based on the scores, fold-changes, and p-values observed in the I-SPY1 and BEAUTY trials.

#### Statistics and Bioinformatics analysis

Differential expression (DE) analysis was performed using the empirical Bayes quasi-likelihood F-test (QLF) to identify genes associated with recurrence for both the pre-NAC and post-NAC data with edgeR^41^. Over representation analysis (ORA) was performed to identify cytobands significantly enriched in the DE genes using WebGestalt^42^. Copy number alterations associated with recurrence were evaluated with Fisher’s exact test. Circos plots were generated with Rcircos^43^. Survival association analysis for the selected biomarkers were independently investigated using the online tool Kaplan-Meier (KM) plotter^44^. In addition, 1) TCGA paired tumor and normal-adjacent TNBC samples^14^, 2) non-LAR vs. LAR TNBC gene expression cohorts 3) non-LAR gene expression cohorts with respect to pathologic complete response phenotype were also surveyed using publically available gene expression data^3,45–49^.

#### Assessment of biomarkers using machine learning methods

The I-SPY1 and BEAUTY studies were pooled together and scaled with the combat algorithm from the sva package (v 3.14.0)^50^ to remove the batch effect. Given the lack of sampling power, we implemented a three-fold cross-validation strategy with the super learner package^51^. We examined six classification algorithms, including a generalized linear model (glmnet), k nearest neighbors (k-NN), neural networks, random forests, support vector machine (SVM), and kernel SVM. The models were implemented using down selection methods (Pearson’s correlation, Spearman’s rank correlation, and random forest). We monitored the area under the receiver operating curve (AUC) to indicate the model’s classification performance.

## Data Availability

All data used in this analysis are publically available. BEAUTY study data are available through dbGap accession ID phs001050.v1.p1. I-SPY1 data are available through the GEO database (GSE32603).

